# Identification of *Escherichia coli* Genes Essential for High-Level, Clinically Relevant, Resistance to Antibiotics

**DOI:** 10.1101/2021.11.10.468170

**Authors:** Esmeralda Z. Reyes-Fernández, Noemie Alon Cudkowicz, Sonia Steiner-Mordoch, Shimon Schuldiner

## Abstract

Antibiotic resistance is one of the biggest public health challenges of our time. Here we present a novel approach to recognizing molecular mechanisms essential for maintaining high-level clinically relevant antibiotic resistance. To identify essential genes in this context, we subjected *Escherichia coli* EV18, a strain highly resistant to quinolones, to random transposon mutagenesis. This strain’s unique advantage is that the screen is performed at very high concentrations of the antibiotic, non-permissive for most strains. The transposon’s insertion affected the transcription of five genes required for the maintenance of resistance in EV18. Three of these genes (*YihO, YhdP*, and *WaaY*) have not been previously identified as essential for high-level antimicrobial resistance (AMR). Our work provides a new perspective to identify and characterize novel players crucial for maintaining AMR in *E. coli*.

## INTRODUCTION

Antibiotic resistance poses a formidable problem in hospital-acquired infections. Many findings suggest that inadequate use of antimicrobials may lead to resistance in bacteria (AMR) and make the treatment of bacterial infections more difficult (1-4). Extensive knowledge of the molecular mechanisms underlying microbial antibiotic resistance is required to fight the increasing numbers of drug-resistant and multidrug-resistant bacteria successfully.

Many genes are involved in *Escherichia coli* and other organisms’ response to antibiotic exposure and AMR acquisition (5). Resistance to antibiotics can be caused generally by the inactivation or modification of the antibiotic, alteration in the target site of the antibiotic, reduction of the intracellular antibiotic accumulation by increasing the active efflux of the antibiotic and modification of metabolic pathways to circumvent the antibiotic effect (6-11).

In an in-lab evolution experiment, we identified Multidrug Transporters (MDTs)’ that play a central role in the process of acquisition of high level clinically relevant resistance but, in this strain, they are not essential for maintenance (12). Therefore, *E. coli* EV18, our evolved strain, provides an excellent tool to identify genes essential for maintenance other than MDTs.

Random mutagenesis has been considered one of the strategies to study genes’ expression and function with crucial roles in different biological processes. Transposition based strategies have been used successfully to generate mutant transposon libraries, for site-specific tagging, and the generation of transcriptional/translational target gene fusions (13-18). To identify genes essential for maintaining high-level, clinically relevant antibiotic resistance, we subjected EV18, to random transposon mutagenesis. The unique advantage provided by this strain is that the screen is performed at very high concentrations of the antibiotic, non-permissive for most strains. We screened ∼4000 colonies for candidate genes whose disruption led to an increase in norfloxacin susceptibility. The insertion of the transposon affected the transcription of five genes required for the maintenance of resistance in EV18, including *acrAB* (RND tripartite transporter), *waaY* (lipopolysaccharide core heptose II kinase), *yihO* (sulfoquinovose transporter), *yhdP* (outer membrane permeability factor), and *emrK* (tripartite efflux pump membrane fusion protein). Three of these genes (*YihO, YhdP*, and *WaaY*) have not been previously identified as essential for high-level AMR. Our work provides a new perspective to identify and characterize novel players crucial for maintaining AMR in *E. coli*.

## RESULTS AND DISCUSSION

### 1. Isolation of *E. coli* EV18-mutants with increased susceptibility to norfloxacin after Tn5 mutagenesis

In a previous study, we described the generation of a highly resistant strain to norfloxacin, EV18, by exposing *E. coli* cells to consecutive increasing concentrations of this antibiotic in an *in-vitro* evolution experiment (12). In that work, we were able to show the role of multidrug transporters (MDTs) and suppliers on the acquisition of norfloxacin-resistance. However, once resistance is achieved, the role of MDTs in maintenance is minor, if at all (12). To understand this phenomenon’s complexity, we performed transposon mutagenesis of the EV18 strain to seek for mutants with a decreased resistance to norfloxacin (**Fig. 1A**). As a first step, we transformed the EV18 cells with the EZ-Tn5 <KAN-2> Tnp Transposome (Epicentre) to create a library (∼ 4000 colonies). In the library’s initial screening we streaked all the colonies resistant to Kanamycin on LB agar plates containing 100 and 400 μM norfloxacin. Mutants showing an increased susceptibility to norfloxacin were isolated and re-tested by growing them in LB liquid medium with norfloxacin at the same concentrations (**Fig. 1A**). We selected only the mutants that displayed sensitivity to norfloxacin on agar and liquid medium for further transposon screening. In total, 52 candidates completed these requirements. For the mapping, the candidates’ genomic DNA was digested to produce blunt-ended fragments and, consequently, ligated to the anchor bubble unit. All the ligations were further amplified by PCR with specific primers and sequenced. We could map up to 50 out of the 52 transposition events. We then tested plasmid-encoded genes’ ability to restore partially or entirely the resistance to norfloxacin and found five that we characterize further, including EV18 Tn5-13 (*acrR-acrA*), EV18 Tn5-19 (*waaY*), EV18 Tn5-32 (*yihO*), EV18 Tn5-47 (*yhdP*), and EV18 Tn5-48 (*emrK*). Once the point of insertion was identified, we generated the corresponding nil strains and all the studies described below, except for Tn5-13, were performed in the nil strains.

**Fig. 1.**
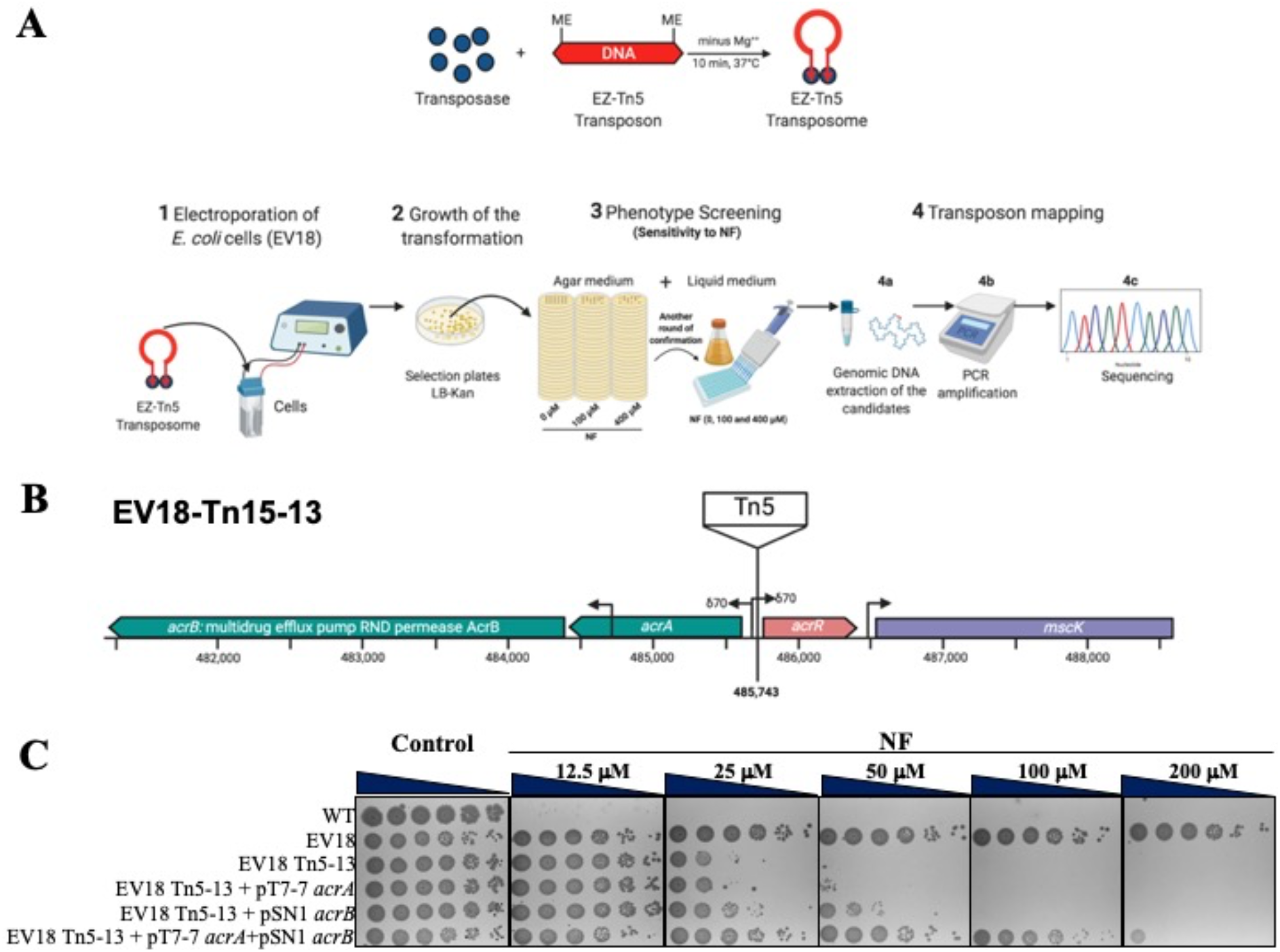
Transposon mutagenesis for identifying candidate genes required for the maintenance of the NF-resistance in the EV18 strain. **(A)** Generation of the transposon library in the NF-resistant *E. coli* strain EV18. **Top:** Formation of the transposon complex EZ-Tn5 by incubating the EZ-Tn5 transposase enzyme with the EZ-Tn5 transposon, containing the Tn903 kanamycin resistance gene (Kan^R^) flanked by the Mosaic End (ME) transposase recognition sequences. **Bottom:** Steps describing the process of selecting EV18-transposition mutants with increased sensitivity to NF. (1) Electroporation of the EV18 cells with the EZ-Tn5 transposome. After recovery of the transformed cells, (2) cells were spread onto LB-Kan agar plates and incubated overnight at (37°C). Each colony (∼ 4000) was picked and streaked on LB-agar plates containing 100 and 400 µM NF to screen for this antibiotic’s sensitivity (3). The growth of selected candidates in LB medium with NF confirmed their sensitivity, which was studied further. (4) The specific Tn5 insertion sites of these mutants were identified by DNA sequencing (for more details, see Materials and Methods). **(B)** Location of the *acrA* and *acrR* genes in the genomic context of *E. coli* (**C)** The sensitivity of the transposon mutant EV18 Tn5-13 to NF. Growth of the EV18-Tn5-13 mutant on LB-agar plates (+/-NF, at 37°C) with or without the complementation plasmids for overexpressing either *acrA* (pT7-7 *acrA*), *acrB* (pSN1 *acrB*) or both. Major restoration of resistance is achieved when *acrA* and *acrB* are simultaneously overexpressed.

### 2. The non-coding region between *acrR* and *acrA* is mutated in EV18 Tn5-13

The EV18 Tn5-13 mutant sequence analysis revealed that the transposon insertion occurred in the non-coding region between *acrR* and *acrA* (**Fig. 1B**). AcrR regulates the expression of genes involved in multidrug transport (19) and also acts as a global repressor for the *mar-sox-rob* regulon (20). This mutant’s resistance phenotype could be rescued to almost full EV18 levels by simultaneously overexpressing *acrA* and *acrB* genes but not by the individual genes (**Fig. 1C**). As depicted in **Fig. 1C**, the EV18 Tn5-13 strain cannot grow with norfloxacin concentrations higher than 25 µM. The plasmid-encoded *acrB* (pSN1 *acrB*) but not *acrA* (pT7-7 *acrA*) confers a small but distinct restoration of resistance. Almost full restoration is observed when both genes *acrA* and *acrB* are overexpressed together (**Fig. 1C**). A similar picture was obtained when the mutant and overexpressing strains were grown in LB liquid medium with increasing norfloxacin (**Fig. S1**).

Although the resistance of EV18 Tn5-13 decreased significantly by at least one order of magnitude it is still more than 10 fold higher than the naïve strain, BW25113. These results confirm our previous analysis of the EV18 Δ*acrB* phenotype where we showed that the medium level resistance is most likely due to the overexpression of other TolC-dependent transporters, mainly AcrF (12).

These results highlight the well-known prominent role of the AcrAB transporter operon in AMR and provide proof of concept for the strategy presented here.

### 3. *emrK* is essential to maintain the high-level resistance of the EV18 strain

The EV18 Tn5-48 mutant sequence showed a disruption in the transcription of *emrK*, coding for a component of the tripartite efflux pump EmrKY-TolC (21), which maps between the *emrY* and *evgL* genes (**Fig. 2A, Top**). We generated an EmrK knockout (EV18 Δ*emrK*) and, as shown in (**Fig. 2B and C**) it displays reduced resistance to norfloxacin, with weak growth at 12.5 µM norfloxacin. Overexpression of *emrK* in the EV18 Δ*emrK* strain (EV18 Δ*emrK* + pCA24N *emrK*) (**Fig. 2B**) fully restored resistance, indicating that the deletion of *emrK* solely caused the growth defect. The finding that the gene’s deletion in the wild strain does not affect resistance (**Fig. 2D**) highlights the advantage of using the EV18 strain to reveal genes’ function in maintaining resistance. EV18 provides a highly dynamic range of antibiotic concentrations that cannot be reached in the wild type. EmrK role in maintenance of resistance to norfloxacin becomes apparent only at high norfloxacin concentrations.

**Fig. 2.**
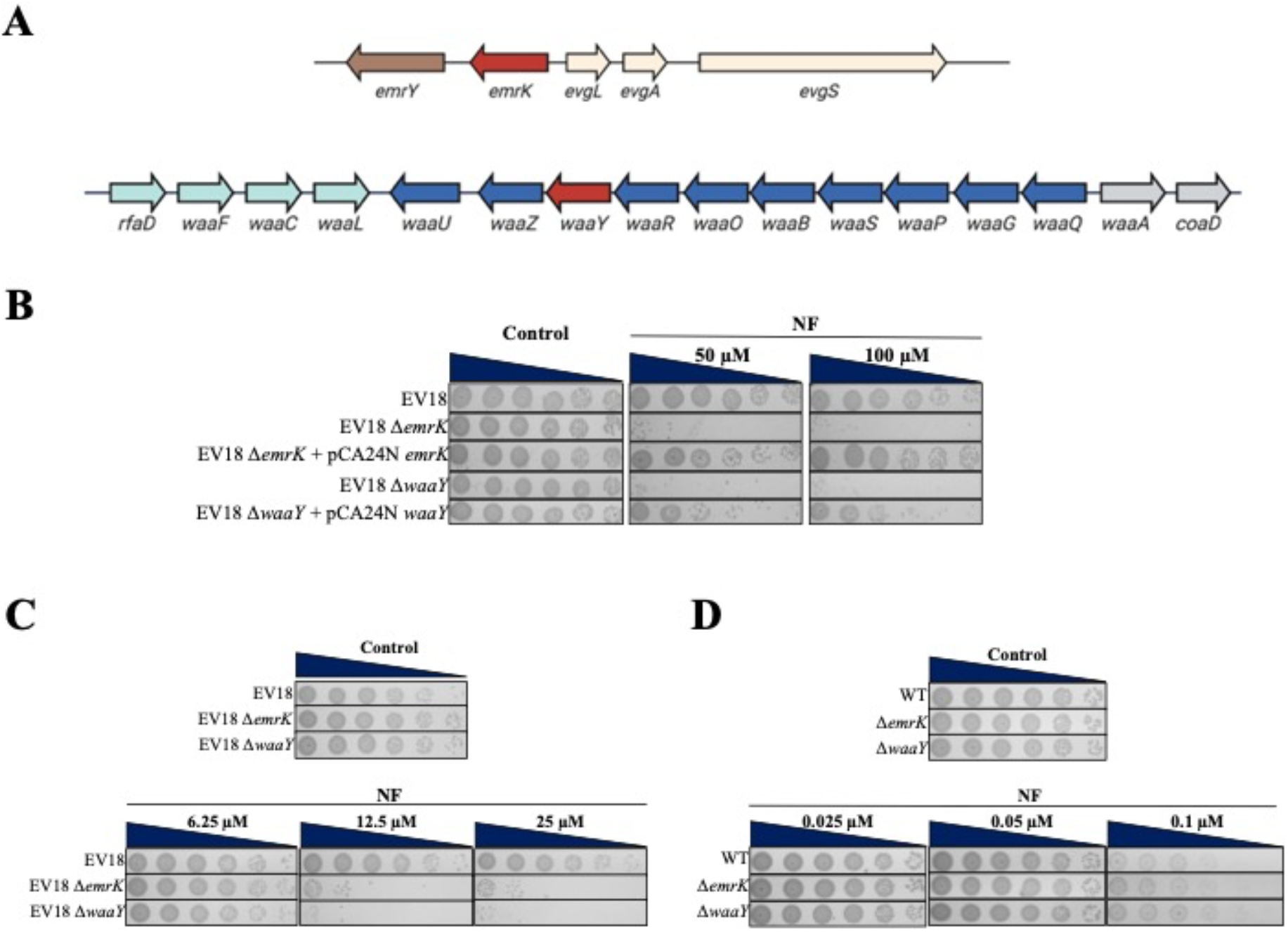
*emrK* and *waaY* are required to maintain the high-level resistance of EV18 to NF. **(A)** Location of the *emrK* and *waaY* genes in the genomic context of *E. coli*. **Top:** The *emrK* gene, encoding the membrane fusion protein of a tripartite efflux pump complex, is flanked by *emrY* and *evgL*, encoding a subunit of the tripartite efflux pump membrane and a small unannotated protein, respectively. **Bottom:** The *waa* locus (formerly *rfa*) contains the three major gene operons for lipopolysaccharides (LPS) biosynthesis in *E. coli*. The operons are identified by the first gene of the transcriptional unit as *rfaD* (green), *waaQ* (blue), and *waaA* (gray). The *waaY* gene, encoding a lipopolysaccharide core heptose (II) kinase, belongs to the *waaQ* operon, which encodes enzymes involved in the core oligosaccharide (OS) assembly. Genes are represented as arrows. **(B-C)** Phenotypes of the Δ*emrK* and Δ*waaY* strains grown on LB-agar plates in the presence of increasing concentrations of norfloxacin (NF), as indicated on top of the plates. Deletion mutants generated in the background of the EV18 strain are shown in **B** and **C**, while the mutants in the WT (BW25113) are in **D**. The growth of the EV18 Δ*emrK* and EV18 Δ*waaY* strains on LB plates with NF was restored almost to the EV18 levels after transformation of the deletion mutants with the plasmids pCA24N *emr*K and pCA24N *waaY* respectively.

To explore the scope of EmrK’s role in supporting antibiotic resistance, we assayed the growth of EV18 Δ*emrK* on agar plates with increasing concentrations of Erythromycin (Ery) and chloramphenicol (Cam) (**Fig. S2A**). The increased sensitivity of EV18 Δ*emrK* to both antibiotics show that the cross-resistance gained by EV18 to other antibiotics, as norfloxacin and chloramphenicol, is compromised when *emrK* is deleted. Interestingly, under the conditions tested, the function of *emrK* appears to be irrelevant in the WT background since its deletion did not generate any change in the phenotype of strain when grown in norfloxacin’s presence (**Fig. 2D**), erythromycin and chloramphenicol (**Fig. S2B**).

The above-described data show that the expression of *emrK* in the EV18 strain is required to preserve its resistance also to Erythromycin and Chloramphenicol.

### 4. Disruption of *waaY* transcription weakens the high-level resistance of the EV18 strain

The DNA sequence of EV18 Tn5-19 revealed that the transposon insertion affected the *waaY* gene’s transcription. In *E. coli*, the *waaY* gene is located in the *waa* operon, a long transcription unit involved in the lipopolysaccharide (LPS) biosynthesis (**Fig. 2A, Bottom**). *waaY* encodes an LPS core heptose (II) kinase that adds a phosphate group to the second heptose residue in the inner core of LPS (22). Changes in the LPS structure can lead to changes in the function and stability of Gram-negative bacteria’s outer membrane (23). Growth of EV18 Δ*waaY* on agar plates revealed a significant decrease in resistance compared to the parent strain, and growth was barely detected at 12.5 µM norfloxacin (**Fig. 2C**). Overexpression of *waaY* restores norfloxacin’s resistance phenotype almost to the EV18 levels: growth is detectable at 100 µM norfloxacin (**Fig. 2B**).

Similar to what we described above for the *emrK* gene, also the cross resistance of EV18 to Erythromycin and chloramphenicol was compromised in the EV18 Δ*waaY* mutant (**Fig. S2A**). Likewise, under the conditions tested, the function of *waaY* appears to be irrelevant in the WT background since its deletion did not generate any change in the phenotype of strain when grown in norfloxacin’s presence (**Fig. 2D**), erythromycin and chloramphenicol (**Fig. S2B**) meaning that the function of *waaY* is not required for the resistance of the WT cells at the antibiotic concentrations tested.

The results point to an essential link between phosphorylation of LPS and the maintenance of antibiotic resistance in the EV18 strain at the high antibiotic concentration.

### 5. Deleting the outer membrane permeability factor encoding gene-*yhdP* causes a decrease in antibiotic resistance in the EV18 strain

Gram-negative bacteria possess a thin peptidoglycan cell wall, surrounded by an outer membrane (OM) containing LPS. The OM is essential for *E. coli* since it acts as a protective barrier against toxic compounds and antibiotics (23). The reduced resistance to norfloxacin of the mutant EV18-Tn5-47 is due to the disruption of the OM permeability factor encoding gene-*yhdP. yhdP* is located between the metalloprotease subunit (*tldD*) and the RNase G (*rng*)-encoding genes (**Fig. 3A, Top**). Previous reports indicate that the deletion of *yhdP* increased the OM’s permeability through interaction with the cyclic enterobacterial common antigen (cyclic ECA) (24). Furthermore, Δ*yhdP* mutants displayed a decrease in SDS and vancomycin resistance (25) and YhdP is involved in modulating the high-flux phospholipid transport pathway to the outer membrane (26).

**Fig. 3.**
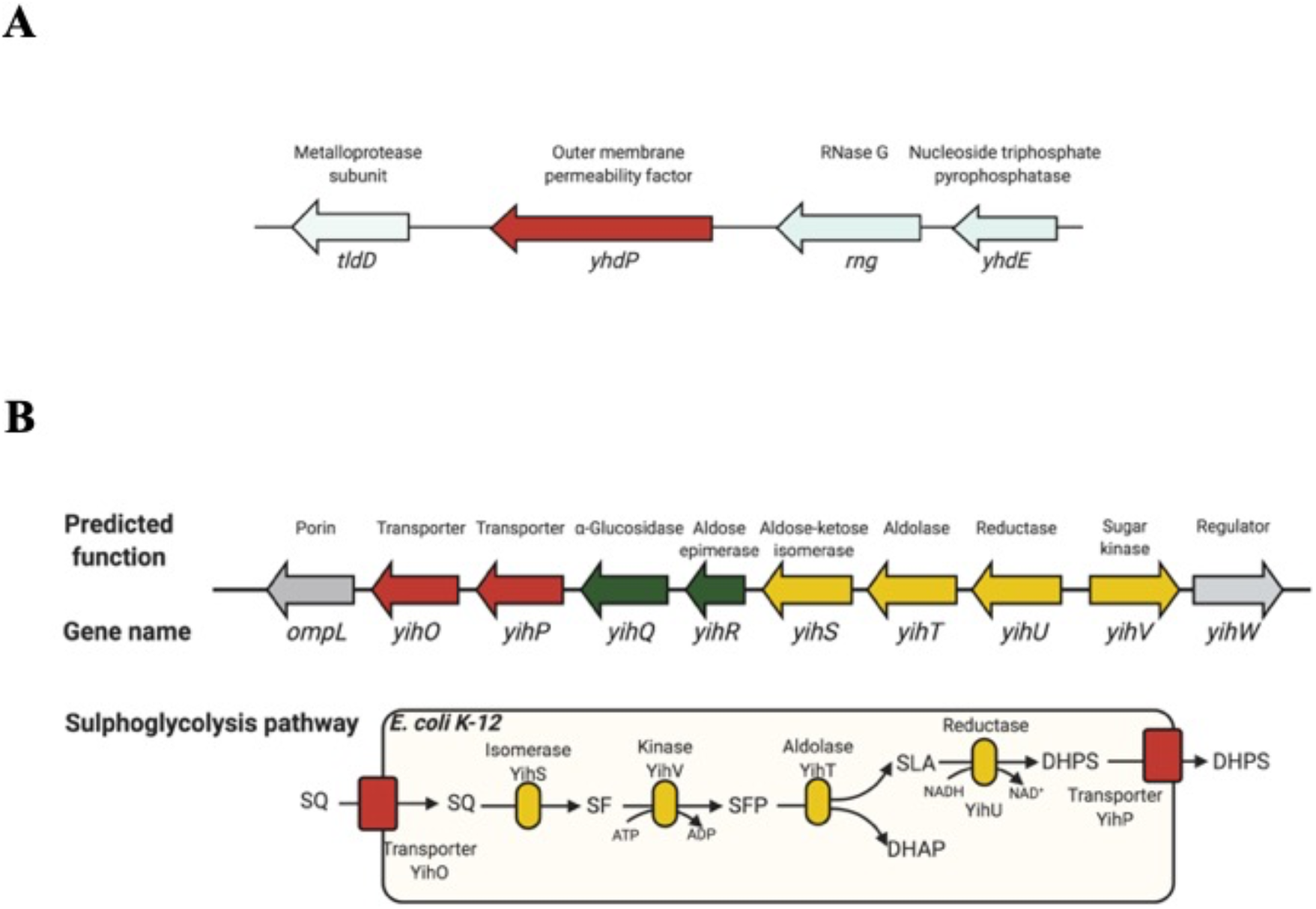
Genomic context of the *yhdP* and *yihO* genes in *E. coli*. **(A)** The *yhdP* gene (in red) encodes for a large protein implicated in maintaining outer membrane permeability through its functional interaction with cyclic enterobacterial common antigen (cyclic ECA). **(B)** Representation of the operon implicated in the sulfoglycolysis pathway. The yihW gene product controls the transcription of the operon. **(Bottom)** Predicted pathway for the catabolism and transport of sulfoquinovose (SQ) (adapted from [Denger, 2014 #8209]). Abbreviations: SQ, sulfoquinovose; SF, 6-deoxy-6-sulfofructose; SFP, 6-deoxy-6-sulfofructose-1-phosphate; SLA, 3-sulfoacetaldehyde; DHAP, dihydroxyacetone phosphate; DHPS, 2,3-dihydroxypropane sulfonate.

As seen in **Fig. 4**, EV18 Δ*yhdP* cells display increased sensitivity to norfloxacin, showing weak growth on agar plates at 25 µM norfloxacin and higher. Overexpression of *yhdP* (pKK *yhdP*) fully reverts the resistance phenotype (**Fig. 4 A**). Thus, we concluded that the YhdP activity’s disturbance causes the sensitivity of EV18 Δ*yhdP* to norfloxacin. Likewise, the cross-resistance of EV18 to norfloxacin and chloramphenicol significantly decreased upon the deletion of *yhdP* (**Fig. 4B**). We detected similar results following the growth of these strains in LB-liquid medium with norfloxacin (25-400 µM), norfloxacin (25-400 µM), and chloramphenicol (0.625-10 µM) (**Fig. S3**). The mutation’s pleiotropic effect is in accordance with the predicted change in OM structure in this mutant, leading to increased permeability of the antibiotics.

**Fig. 4.**
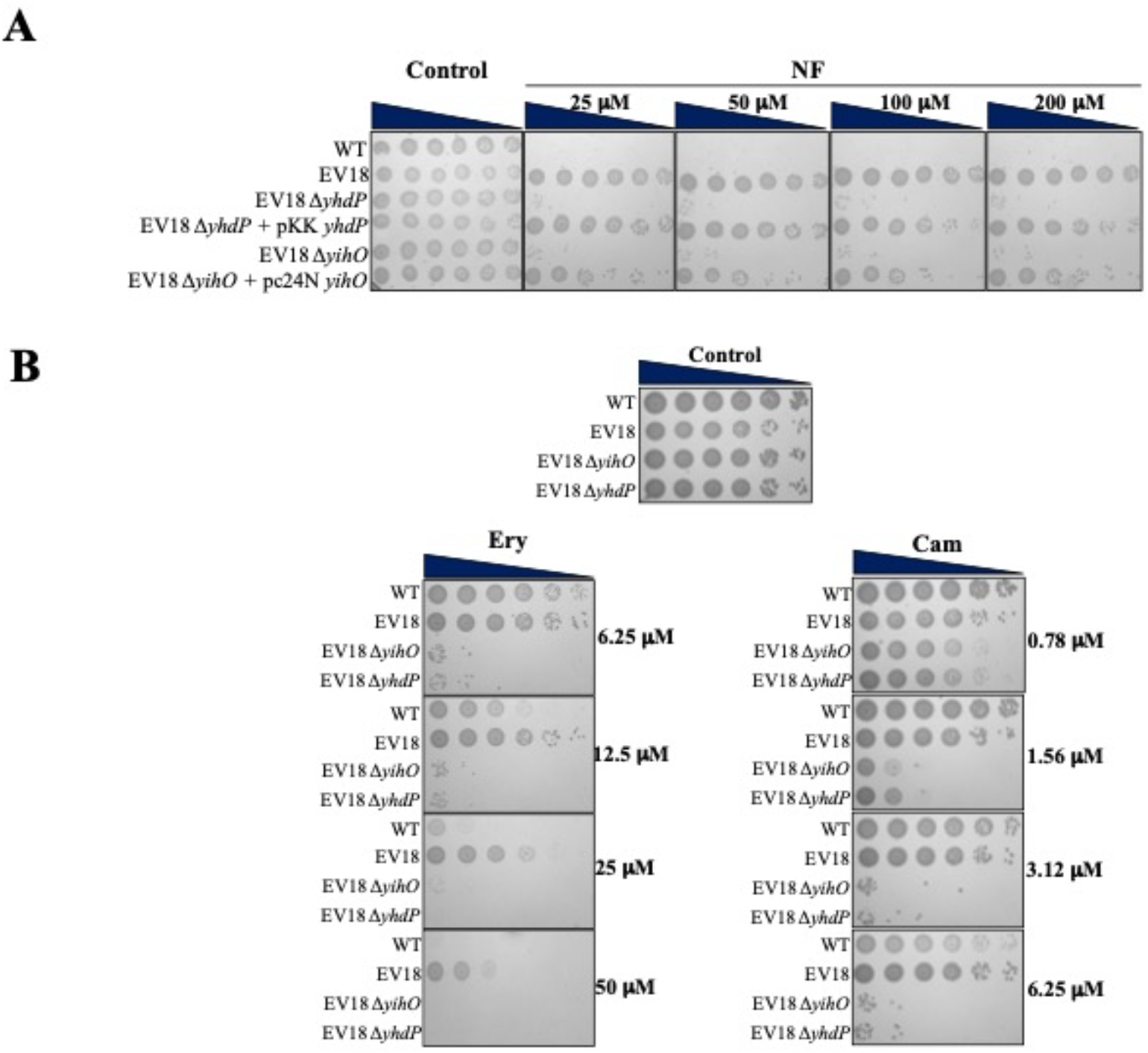
*YihO* and *yhdP* are required to maintain the high-level resistance of EV18 to NF, Ery, and Cam. **(A)** The deletion of *yihO* and *yhdP* reduces the resistance of the EV18 strain to NF. Quantitative restoration of the resistance is achieved in cells bearing plasmids pC24N *yihO* and pKK *yhdP*, respectively. Concentrations of NF are indicated on top of each plate. All plates included 50 µM IPTG. **(B)** The deletion of *yihO* and *yhdP* also reduces the resistance of the EV18 strain to Ery and Cam (left and right, respectively). Concentrations for each antibiotic are indicated on the right side of the plates.

### 6. A transporter of the sulfoglycolysis pathway, YihO, is required to maintain the antibiotic resistance in the EV18 strain

Sulfoquinovose (SQ; 6-deoxy-6-sulfoglucose) is one of the most abundant compounds in nature and the principal component of plant sulfolipid (27). *E. coli* K-12 can use SQ as a sole carbon source and energy for growth in the so-called sulfoglycolysis (SFG) pathway **(Fig 3B)** (28). In this pathway, SQ is transported by the YihO transporter. Afterward, SQ is isomerized to sulfofructose (SF) by the SQ-isomerase YihS. Then, SF is phosphorylated to SF-6-phosphate (SFP), a step catalyzed by the SF-kinase YihV. SFP is subsequently cleaved by the SFP-aldolase (YihT) into sulfolactaldehyde (SLA) and dihydroxyacetone phosphate (DHAP). Finally, SLA is reduced to 2,3-dihydroxypropane-1-sulfonate (DHPS) by the SLA-reductase YihU. Another transporter in the operon, YihP, is 65% identical in sequence to YihO. It may function to export DHPS but it has not been experimentally characterized. Besides the genes required for SQ transport and catabolism, an additional gene, *yihW* encoding a transcription factor, is located downstream of the *yihV* gene. A representation of this operon and the suggested sulfoglycolysis pathway is seen in **Fig. 3B**. In our work, we identified a mutant (EV18-Tn5-32) whose *yihO*-transcription was affected by the transposon’s insertion in the coding region of this gene. By deleting *yihO* in the EV18 strain, we noticed a remarkable decrease in antibiotic resistance, not only to norfloxacin (**Fig. 4A**), but also to erythromycin, and chloramphenicol on agar plates (**Fig. 4B**). The rescue of the resistance phenotype by overexpressing *yihO* (pCA24N *yihO*) confirmed the relevance of *yihO* in the maintenance of resistance of EV18 **(Fig. 4A)**. As previously described for the *emrK* and *waaY* knockouts, the sensitivity of WT Δ*yihO* to the above mentioned antibiotics did not increase **(Fig S4)**.

Since *yihO* codes for the first protein in the sulfoglycolysis pathway (**Fig. 3B**), we also constructed the knockout mutants of the genes encoding proteins downstream of *yihO* in the pathway, a putative 2,3-dihydroxypropane-1-sulfonate export protein (*yihP*), a 6-deoxy-6-sulfofructose-1-phosphate aldolase (*yihT*), and a 3-sulfolactaldehyde reductase (*yihU*). The results reveal that the resistance to norfloxacin, erythromycin, and chloramphenicol remains unmodified compared to EV18 (**Fig. 5**). In other words, EV18 Δ*yihO* was the only mutant from the SQ operon (analyzed in this study) whose resistance to norfloxacin, erythromycin, and chloramphenicol underwent a substantial decrease.

**Fig. 5.**
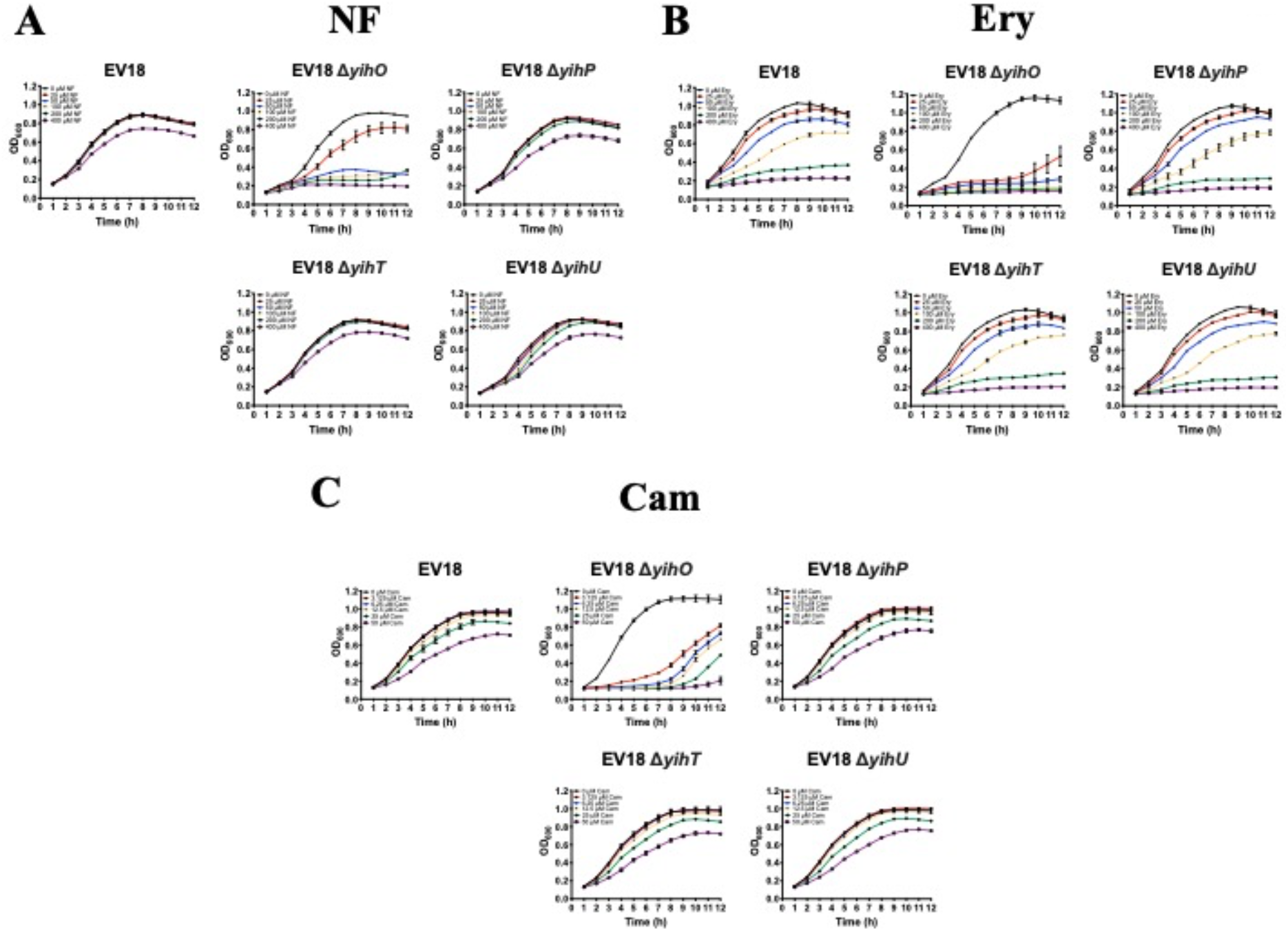
The deletion of *yihO* but not of other genes in the operon affects the resistance of the EV18 strain to several antibiotics. Liquid cultures of EV18, EV18 Δ*yihO*, EV18 Δ*yihP*, EV18 Δ*yihT*, and EV18 Δ*yihU*, were prepared in LB medium to an OD_600_=0.01 and transferred to a microtiter plate with the indicated concentrations of antibiotics. The growth of the cells was recorded every hour by a microtiter plate reader. The antibiotics used in this study were **(A)** NF, **(B)** Ery, and **(C)** Cam. From all the deletion mutants, only EV18 Δ*yihO* showed an effect on growth with all the tested antibiotics.

*yihO* codes for a member of the glycoside-pentoside-hexuronide (GPH) cation symporter family of transporters (29). Although its transport mechanism has not been experimentally characterized it has been suggested that it functions as a SQ transporter (28). Since deletion of other genes in the sulfoglycolysis pathway does not have an effect on EV18 resistance we hypothesize that YihO may have other functions essential for resistance.

Therefore, to provide a clue to the possible role of YihO in EV18 resistance, we screened for potential substrates, other than SG. LB is a commonly used nutritionally rich medium for culturing bacteria with a complex composition. To test whether the effect of the deletion of *yihO* is also observed in media with defined composition, we tested the growth of the WT Δ*yihO* and EV18 Δ*yihO* strains on MMA agar plates with norfloxacin in the presence of different carbon sources, including glucose, glycerol, maltose, and sodium succinate (**Fig. S5**). In all cases, the growth of EV18 Δ*yihO* was undetectable at 25 µM norfloxacin **(Fig. S5B)**. As shown before for the LB medium, under the conditions tested, the deletion of *yihO* does not render the WT strain more sensitive to norfloxacin (**Fig. S5A**). We conclude that the effect of the *yihO* deletion on the phenotype is detected in rich medium as well as in the defined media with various carbon sources (**Fig. S5B**).

To check whether YihO also works as a drug exporter, we tested the possible impact of YihO overexpression in wild-type cells on resistance to several antibiotics and toxicants but detected no effect **(Table S1)**.

A closer look at the SQ operon’s genomic location points to the downstream proximity of a putative outer membrane porin encoded by the gene *ompL* **(Fig. 3B)**. OmpL localizes to the outer membrane and exhibits porin-type properties allowing solutes smaller than 600 Daltons to pass into and out of the periplasm (30). To rule out the possibility that the deletion of *yihO* influenced the expression of *ompL*, we tested the resistance of EV18 Δ*ompL* to different antibiotics **(Fig. S6)**. The growth of EV18 and EV18 Δ*ompL* revealed that *ompL* deletion does not alter the resistance of EV18 to norfloxacin, norfloxacin, and chloramphenicol **(Fig. S6)**.

We conclude that the loss of resistance of EV18 Δ*yihO* is due to the *yihO* deletion and rules out a possible positional effect on the expression of *ompL*. Moreover, the effect of the deletion is not due to an alteration in the sulfoglycolysis pathway and further work is required to elucidate the mechanistic basis of this finding.

## CONCLUSION

To fight the increasing numbers of drug-resistant and multidrug-resistant bacteria successfully, we need a detailed knowledge of the molecular mechanisms underlying the acquisition and maintenance of microbial antibiotic resistance.

The approach described in this paper offers an unbiased effort that allows the identification of genes essential for maintaining the very high-level clinically-relevant resistance. Random transposon mutagenesis has been extensively and successfully used prior to this publication. The unique advantage of the work described here is that thanks to the isolation in our lab of the E. coli strain EV18, screening of mutants with decreased resistance were performed at very high concentrations of antibiotics, in our case, norfloxacin.

## MATERIALS AND METHODS

### 1. Bacterial strains, growth media, and culture conditions

Bacterial strains, plasmids and primers used in this study are shown in **Tables S2-S4**. *Escherichia coli* strains BW25113 were grown in LB-Na broth (10g/L tryptone, 5g/L yeast extract and 5g/L NaCl), LB-NaPi broth (LB-Na broth with 70 mM sodium phosphate, pH=7.4) or minimal medium A (MMA)(31) with 0.5% glycerol, glucose, maltose or sodium succinate as a carbon source, and supplemented with 1X MEM Amino Acids solutions M550 and M7145. All the antibiotics and reagents used in this study were purchased from Sigma-Aldrich, except for ampicillin (Amp), was from Teva Pharmaceuticals (Petach Tikva, Israel). All the deletion mutants analyzed in this study were created as described by Datsenko and Wanner (32).

### 2. Transposon insertion mutagenesis

EV18 strain was used as the recipient strain for transposon mutagenesis and gene construction. Transposon insertion into *E. coli* chromosome was carried out by electroporating 1 μL of the EZ-Tn5 <KAN-2> Tnp Transposome (Epicentre) in 50 μL of EV18-pkD46 electrocompetent cells, followed by 2 h recovery in 1mL of SOC medium [0.5% yeast extract, 2% tryptone, 10 mM NaCl, 2.5 mM KCl, 10 mM MgCl_2_, 10 mM MgSO_4_, and 20 mM glucose] at 37°C. Afterwards, the whole transformation was plated on Kan-containing plates (100 μL/plate), and incubated overnight at 37°C. To identify sensitive clones, about 4000 colonies were picked and transferred to LB-norfloxacin plates (0, 100 and 400 μM). To confirm their sensitivity to norfloxacin, colonies that did not grow on 400 and/or 100 μM norfloxacin were subjected to another round of analysis in LB liquid medium. Finally, colonies sensitive to norfloxacin on agar and in liquid medium were subjected to transposon mapping.

### 3. Transposition mapping

Transposition mapping was done essentially as described in (33). Genomic DNA was prepared using a NucleoSpin kit (Macherey-Nagel) according to the manufacturers’ instructions. Subsequently, gDNA was digested overnight (37°C) with the restriction enzyme RsaI (New England Biolabs) to produce small, blunt-ended fragments. After digestion, 2 μL of anchor bubble unit (see the preparation below), 1 μL of 10 mM ATP and 1 μL of T4 DNA ligase (New England Biolabs) were added and the ligation reaction was incubated for 5 cycles at 20°C for one hour followed by 37°C for 30 min. The anchor bubble unit was prepared by annealing the bubble primers (**Table S3**) in equal amount (4 μM of each primer) in a total volume of 100 μL. The mixture was incubated for 5 minutes at 65°C, and then MgCl_2_ to a final concentration of 1-2 mM before cooling down to room temperature. Following ligation, PCR amplification was performed using 2μL of DNA templates (RsaI-digested DNA ligated to the bubbles, as prepared above), 1 μL of 20 mM of each primer (KAN-2 FP1 or KAN-2 RP1 complementary to the Tn5 sequence with the bubble primer 224) and 10 μL of the 10X Qiagen Multiplex PCR Master Mix Kit (Qiagen, Valencia, CA, USA) in a final volume of 100 μL. PCR cycling conditions were: 95°C for 15 min, 35 cycles of 94°C for 30 s, 60°C for 90 s, 72°C for 2 min, and a final extension step at 72°C for 10 min. The amplified PCR products were separated in 1% agarose, stained with ethidium bromide and visualized under UV light. DNA bands were excised, purified with the PureLink Qiagen DNA Gel Extraction Kit and sent out for sequencing. DNA sequencing analysis was carried out using specific primers to the Tn5 and bubble regions. DNA sequences were analyzed by BLAST and compared to the EV18 sequence in order to find the accurate insertion place of the transposon.

### 4. Susceptibility assays to antibiotics of *E. coli* strains

Pre-cultures were grown overnight in 4 mL of LB-Na and diluted 1:100 in either LB-Na or MMA (pH=7.4). Cells continued growing for several hours (until early logarithmic phase) and diluted to a final OD_600_=0.02 in either LB-NaPi or MMA for examining their growth in liquid media. Duplicate two fold serial dilutions were made in 96-well for each antibiotic, and one well only with medium in order to rule out contamination. Growth was followed for 16 h, measuring the optical density at 600 nm every hour, at 37°C and constant shaking in a Synergy 2 Microplate Reader (Bio Tek). Results were obtained by comparing the percentage of growth of the tested strain grown without antibiotics with the strain grown in the presence of them. Growth curves were obtained by using the GraphPad Prism 7 software. For the phenotypes on solid medium, the cells were grown as indicated above but instead of diluting them to an OD600=0.02, they were brought to OD600=0.2 and 5 μL of 10-fold serial dilutions of the cells were spotted on LB-NaPi (pH=7.4) agar plates containing increasing concentrations of antibiotics according to the experimental setup. Plates were then incubated (37°C, overnight), and growth of the strains was recorded by using the Fusion Fx ECL and Gel Documentation System (VILBER). Biological replicates were included for each experiment.

### 5. Sofware

The processing of the graphs was performed with GraphPad Prism 7 software. The images describing the methodology for the transposon mutagenesis (Fig. 1), as well as in Fig. 2 and 3 shown in this work were created with BioRender.

## Abbreviations

(AMR): Antimicrobial Resistance
(MMA): Minimal Medium A
(norfloxacin): Norfloxacin

## Acknowledgment

SS is Mathilda Marks-Kennedy Emeritus Professor of Biochemistry at the Hebrew University of Jerusalem. The authors declare non-competing interests.

