## Supplemental Figures and Tables for "Identification of *Escherichia coli* Genes Essential for High-Level, Clinically Relevant, Resistance to Antibiotics"

By

Esmeralda Z. Reyes-Fernández[1,2], Noemie Alon Cudkowicz[1], Sonia Steiner-Mordoch[1] and Shimon Schuldiner*

**Supplementary Material**

**Fig.S1**

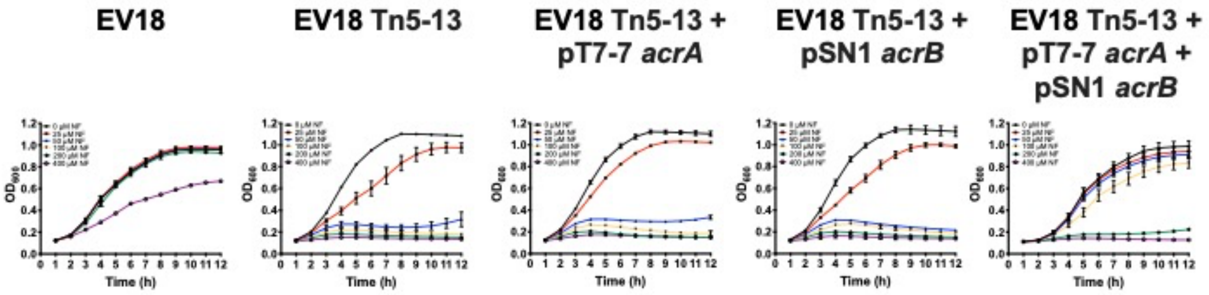


**Fig. S1: Analysis of the growth in LB liquid medium of the strains described in Fig. 1 at increasing concentrations of NF.**

Fig.S2

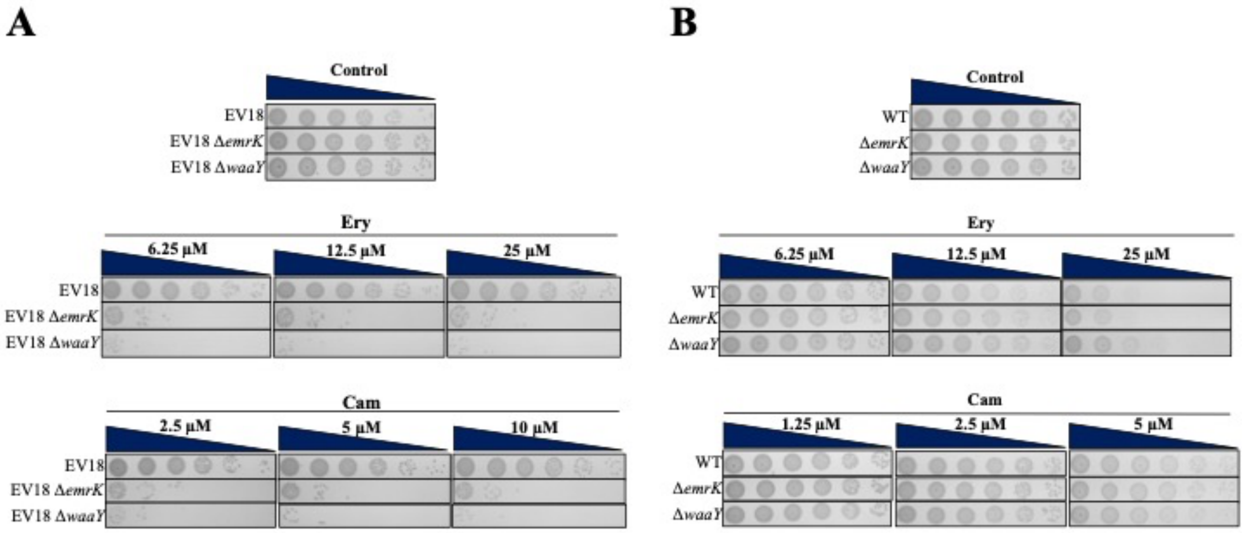


**Fig. S2. Analysis of the sensitivity of the EV18 Δ*emrK* and EV18 Δ*waaY* strains to Erythromycin and Chloramphenicol (A)** The growth of the EV18 Δ*emrK* and EV18 Δ*waaY* strains on LB plates with various concentrations of Erythromycin and Chloramphenicol.

Fig.S3

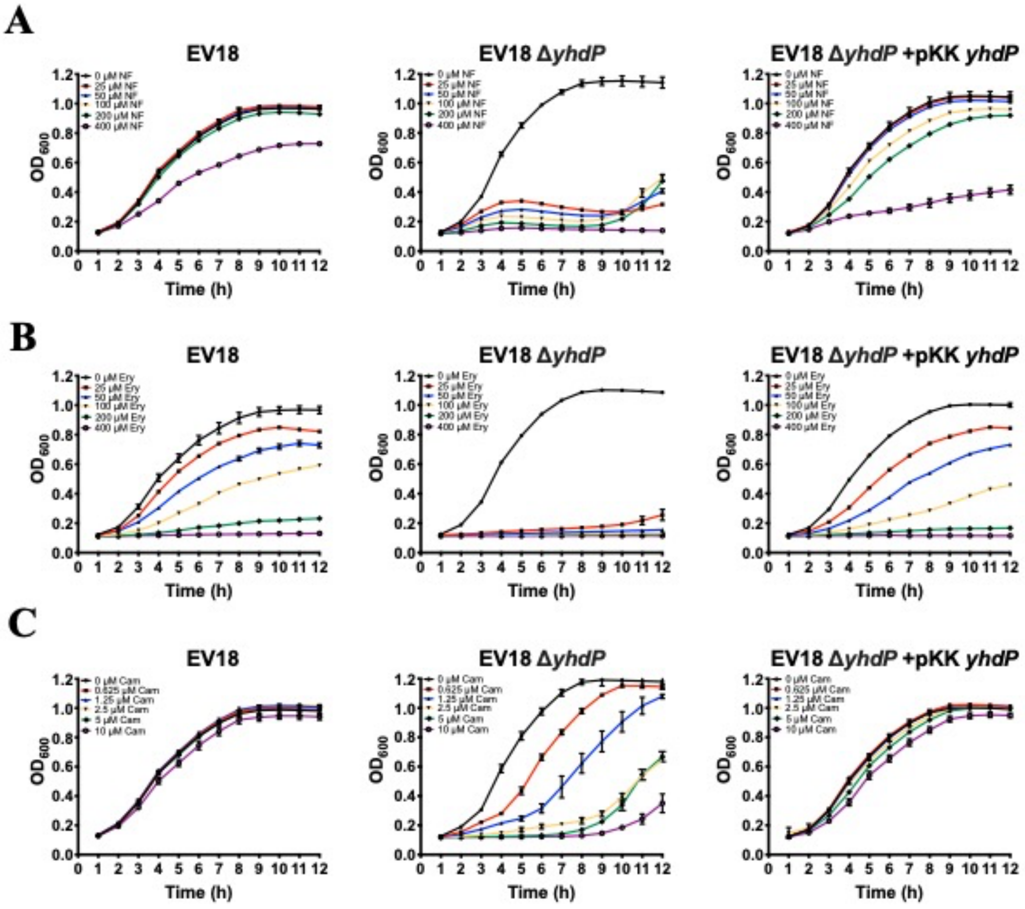


**Fig. S3. The *yhdP* gene has an essential role in the resistance of EV18 to NF, Ery, and Cam.** Analysis of the growth of the EV18, EV18 Δ*yhdP* and EV18 Δ*yhdP* with the complementation plasmid (pKK *yhdP*) was carried out in LB liquid medium for 12 h (37°C) in the presence of either NF **(A)**, Ery **(B)**, or Cam **(C)**. The deletion of *yhdP* renders the EV18 strain more sensitive to NF, Ery, and Cam. Transformation of EV18 Δ*yhdP* with the pKK *yhdP* plasmid restores the strain resistance almost to the EV18 levels.

**Fig.S4**

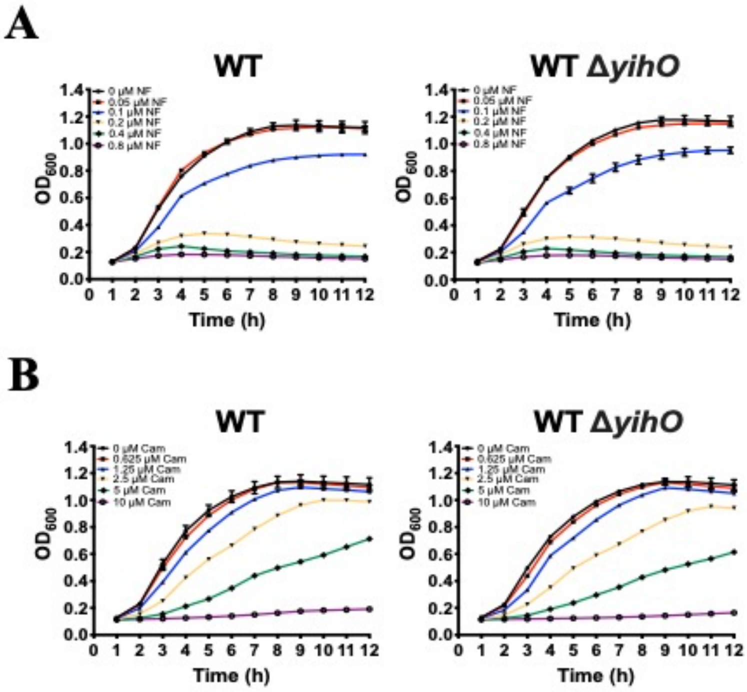


**Fig. S4. Deletion of *ompL* does not impair resistance of the EV18 strain to NF, Ery, Cam and Gen.** The growth of the EV18 and EV18 Δ*ompL* strains was monitored in LB liquid medium for 12 h at 37°C in the presence of different antibiotics including NF **(A)**, Ery **(B)**, and Cam **(C)**. The concentrations for each antibiotic are indicated on the top-left corner of each graph.

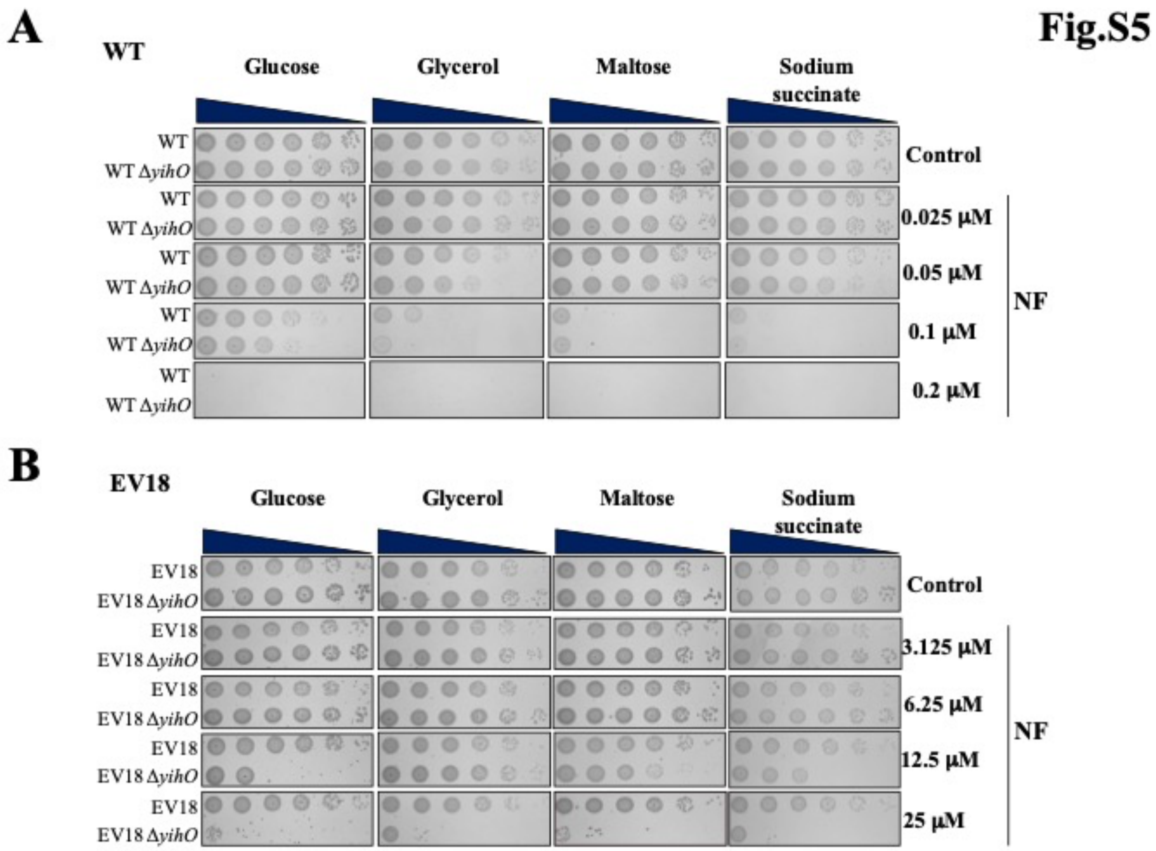


**Fig. S5. The deletion of the *yihO* gene does not affect the WT sensitivity to NF and Cam.** Growth of the WT Δ*yihO* **(A)** and EVC Δ*yihO* **(B)** strains in the presence of increasing concentrations of NF **(Top)** and Cam **(Bottom).** The concentrations of NF and Cam are indicated on the top-left corner for each graph.

Fig.S6

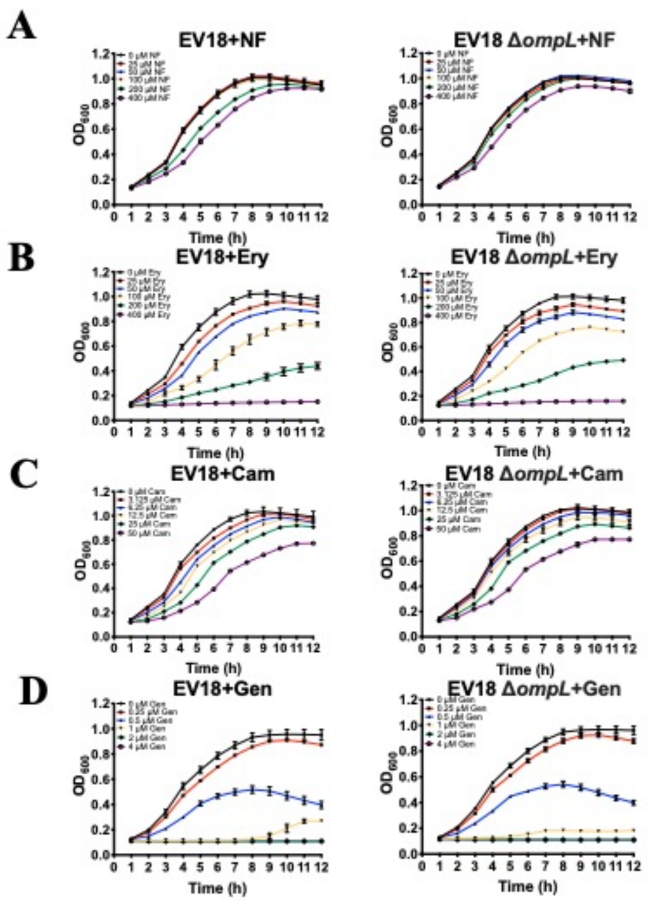


**Fig. S6. The effect of different carbon sources on the sensitivity to NF of the WT Δ*yihO* (A) and EV18 Δ*yihO* (B) strains**. Growth was analyzed by a spot assay on MMA-agar plates prepared with either glucose, glycerol, maltose, or sodium succinate. Concentrations of NF for each case are indicated on the right side of the plates.

**Table S1. Overexpression of YihO does not increase antibiotic resistance**

| | $IC_{50}$ (μM) | |
|---|---|---|
| | WT | WT/pCA24N yihO |
| Norfloxacin | 0.095 ± 0.007 | 0.093 ±0.008 |
| Nalidixic acid | 38.0 ± 3.5 | 35.5 ± 3.8 |
| Kanamycin | 2.3 ±0.3 | 2.1 ±0.3 |
| Acriflavine | 32.2 ± 2.8 | 21.3 ±2.6 |

The growth of indicated strains was monitored in LB liquid medium for 12 h at 37°C in the presence of increasing concentrations of antibiotics.

**Table S2. *E. coli* strains used for this study.**

| **Strain** | **Description** | **Reference** |
|---|---|---|
| WT | Wild-type strain (BW25113) | This study |
| Δ*yihO*::*kan* | Deletion mutant of the *yihO* gene in the background of the WT strain. | This study |
| Δ*yhdP*::*kan* | Deletion mutant of the *yhdP* gene in the background of the WT strain. | This study |
| Δ*waaY*::*kan* | Deletion mutant of the *waaY* gene in the background of the WT strain. | This study |
| Δ*waaQ*::*kan* | Deletion mutant of the *waaQ* gene in the background of the WT strain. | This study |
| Δ*waaO*::*kan* | Deletion mutant of the *waaO* gene in the background of the WT strain. | This study |
| Δ*rfaD*::*kan* | Deletion mutant of the *rfaD* gene in the background of the WT strain. | This study |
| Δ*emrK*::*kan* | Deletion mutant of the *emrK* gene in the background of the WT strain. | This study |
| EV18 | NF resistant strain created by evolution *in vitro* (18 days of habituation). | (1) |
| EV18 Tn5-13 | EV18 mutant with insertion of the transposon in the non-coding region between *acrA* and *acrR*. | This study |
| EV18 Δ*yihO*::*kan* | Deletion mutant of the *yihO* gene in the background of the EV18 strain. | This study |
| EV18 Δ*yihP*::*kan* | Deletion mutant of the *yihP* gene in the background of the EV18 strain. | This study |

| | | |
|---|---|---|
| EV18 Δ*yihT*::*kan* | Deletion mutant of the *yihT* gene in the background of the EV18 strain. | This study |
| EV18 Δ*yihU*::*kan* | Deletion mutant of the *yihU* gene in the background of the EV18 strain. | This study |
| EV18 Δ*ompL*::*kan* | Deletion mutant of the *ompL* gene in the background of the EV18 strain. | This study |
| EV18 Δ*yhdP*::*kan* | Deletion mutant of the *yhdP* gene in the background of the EV18 strain. | This study |
| EV18 Δ*waaY*::*kan* | Deletion mutant of the *waaY* gene in the background of the EV18 strain. | This study |
| EV18 Δ*waaQ*::*kan* | Deletion mutant of the *waaQ* gene in the background of the EV18 strain. | This study |
| EV18 Δ*waaO*::*kan* | Deletion mutant of the *waaO* gene in the background of the EV18 strain. | This study |
| EV18 Δ*rfaD*::*kan* | Deletion mutant of the *rfaD* gene in the background of the EV18 strain. | This study |
| EV18 Δ*emrK*::*kan* | Deletion mutant of the *emrK* gene in the background of the EV18 strain. | This study |
| EVC | Cam resistant strain created by evolution *in vitro* (21 days of habituation). | (2) |
| EVC Δ*yihO*::*kan* | Deletion mutant of the *yihO* gene in the background of the EVC strain. | This study |
| EVC Δ*waaY*::*kan* | Deletion mutant of the *waaY* gene in the background of the EVC strain. | This study |
| EVC Δ*waaQ*::*kan* | Deletion mutant of the *waaQ* gene in the background of the EVC strain. | This study |
| EVC Δ*waaO*::*kan* | Deletion mutant of the *waaO* gene in the background of the EVC strain. | This study |
| EVC Δ*rfaD*::*kan* | Deletion mutant of the *rfaD* gene in the background of the EVC strain. | This study |
| EVC Δ*emrK*::*kan* | Deletion mutant of the *emrK* gene in the background of the EVC strain. | This study |

**Table S3. Plasmids used in this study.**

| **Plasmid** | **Reference** |
|---|---|
| pKD13 | (3) |
| pKD46 | (3) |
| pT7-7 | (4) |

| | |
|---|---|
| pKK223-3 | Pharmacia Biotech. Inc. |
| pSN1 | (5) |
| pT7-7 acrA | (6) |
| pSN1 acrB | (6) |
| pKK yhdP | This study |
| pCA24N yihO | (7) |
| pCA24N waaY | (7) |
| pCA24N emrK | (7) |

**Table S4. Primers used in this study.**

| **Primer** | **Sequence (5´- 3´)** | **Tm (°C)** |
|---|---|---|
| *yihO*_H1P1 | CGTAATAATAATAAACGACGCCCTGCGGGGCGTTAT AAGGAGTGATTATGATTCCGGGGATCCGTCGACC | 70.6 |
| *yihO*_H2P2 | ACCAGGATGGAAATAGCGGCTATTGACATTTATAAC GTTACGGAAGCCGTTGTAGGCTGGAGCTGCTTCG | 71 |
| *yihO*_L1 | CGAGAAGATGTATGTCCGCA | 54.4 |
| *yihO*_L2 | AGATAATAAAATTATTGCAT | 39.8 |
| *yihP*_H1P1 | | |
| *yihP*_H1P1 | | |
| *yihP*_L1 | GTCTTTTATCGCGCCGATAC | 53.5 |
| *yihP*_L2 | CCCATAGCAAAGATAATCCTTG | 51.2 |
| *yihT*_H1P1 | GAATAAGTACACCATCAACGACATTACGCGCGCATC GGGCGGTTTTGCCAATTCCGGGGATCCGTCGACC | 73.2 |
| *yihT*_H2P2 | GCGGCGTTTAGCCATCATTTCGTCGACGATATCGCCA AGTTGTTGTAATTTGTAGGCTGGAGCTGCTTCG | 71.3 |
| *yihT*_L1 | CCGGGATGACTGCCAAAGTA | 57.1 |
| *yihT*_L2 | CAACGGTTGTGGCTTAGGGT | 57.8 |
| *yihU*_H1P1 | CGCCACGTATTTACCGCTCTCCGTCGGTAACCCTTCC ACGTAATAGATGCATTCCGGGGATCCGTCGACC | 72.7 |
| *yihU*_H2P2 | CTTTGAAATCAGTTAAAACGCTATCGGCTACCGGAG CGGGTGCCCCAGCCTGTAGGCTGGAGCTGCTTCG | 73.5 |
| *yihU*_L1 | CGCGACCAATAAAATCGACC | 54.3 |
| *yihU*_L2 | CGCAACTTTTGGCAATCGCG | 58.4 |
| *yhdP*_H1P1 | CGGCAAAGGGTTTTTGAGTCACATTTTTAGCAGACA AGGAGTGACGGGTGATTCCGGGGATCCGTCGACC | 72.2 |
| *yhdP*_H2P2 | GATTGGGGCAATTACGCGCCCTCGTCAAATCATTGC GCTTTTTCTTTACGTGTAGGCTGGAGCTGCTTCG | 72.1 |
| *yhdP*_L1 | AACCAGGAGCAGTTTGACGT | 56.9 |
| *yhdP*_L2 | CGGCGTGAACGCCCTATCCG | 63.3 |
| *ompL*_H1P1 | GCTATTTCCATCCTGGTGGTGGCGGCCTCCCTACGT TTAAAAAATGGACATTCCGGGGATCCGTCGACC | 73.1 |

| | | |
|---|---|---|
| *ompL*_H2P2 | GAGATGCTGGCGGCTGAAGCTTAATTAATGGCCGGA TGTGCTTAATCCGGTGTAGGCTGGAGCTGCTTCG | 72.2 |
| *ompL*_L1 | CTGCATTGACGATGGGATTC | 53.9 |
| *ompL*_L2 | GTGTTTGGCATTATGGCCTG | 54.7 |
| *waaY*_H1P1 | GCAACATCATTATATCTCAGGAATTATAGCAGGAGT CTGTTATCTTTGCCATTCCGGGGATCCGTCGACC | 69.2 |
| *waaY*_H2P2 | AATAAATAACTCCGCTTAAAACTGTCCTGGTGAAGA CATGTAATTGTGGGTGTAGGCTGGAGCTGCTTCG | 69.8 |
| *waaY*_L1 | GACTCTCCACGAGATGCGAA | 56.6 |
| *waaY*_L2 | GCGATGTAGGGCCCGAAAGA | 59.7 |
| *waaQ*_H1P1 | GCGATATCATGGGGATATGTTATTAACTACTCCTGTC ATCAGTACGCTCAATTCCGGGGATCCGTCGACC | 69.8 |
| *waaQ*_H2P2 | CATAATTGTGCATTCCTGTCAGCTGAAGGGGCATCTT CTGGCAACAGCTTTGTAGGCTGGAGCTGCTTCG | 72 |
| *waaQ*_L1 | GAGTCGCTAGTGGAAAAGCC | 55.7 |
| *waaQ*_L2 | CAACTGTTGATGCAATGCGC | 55.8 |
| *waaO*_H1P1 | GACATCGCTTATGGAACTGACAAAAACTTTCTATTTG GTTGCGGAATTTCATTCCGGGGATCCGTCGACC | 70 |
| *waaO*_H2P2 | CCTCCATAAAGGCTTGTGATACAGGATAATCCCAGG CCCAATCATGCCAGTGTAGGCTGGAGCTGCTTCG | 72 |
| *waaO*_L1 | CCTGCGTCGATCGATGAATT | 55.3 |
| *waaO*_L2 | AGGAAATGAGTCCACAATGC | 53 |
| *rfaD*_H1P1 | TGATCATCGTTACCGGCGGCGCGGGCTTTATCGGCA GCAACATCGTTAAAATTCCGGGGATCCGTCGACC | 74 |
| *rfaD*_H2P2 | TATGCGTCGCGATTCAGCCAGGCCATGTATTCCGTTA CACCTTCAGCAACTGTAGGCTGGAGCTGCTTCG | 72.9 |
| *rfaD*_L1 | CGTGTCTGAGATTGTCTCTG | 52.6 |
| *rfaD*_L2 | CATGCAGAGCTCATGTCGCCA | 60.2 |
| *emrK*_H1P1 | CAGGTGCCTATGCCTATTGGTCAATGGAATTAGAAG ACATGATTAGTACAATTCCGGGGATCCGTCGACC | 69.8 |
| *emrK*_H2P2 | CTGGTATAAGCCGGCATGGAGGTCACGGTTGAAGCC AGCTCAGGCATCTCTGTAGGCTGGAGCTGCTTCG | 73.7 |
| *emrK*_L1 | GCACTGTGTTTGATATGAAG | 49.2 |
| *emrK*_L2 | CACCGGTTAATGGTGCCGGA | 60.2 |
| K1 | CAGTCATAGCCGAATAGCCT | 54.1 |
| K2 | CGGTGCCCTGAATGAACTGC | 59.2 |
| KT | CGGCCACAGTCGATGAATCC | 58.4 |
| KAN-2 FP1 | ACCTACAACAAAGCTCTCATCAACC | 59 |
| KAN-2 RP1 | GCAATGTAACATCAGAGATTTTGAG | 55 |
| Bubble_1 | GACTCTCCCTTCTCGAATCGTAACCGTTCGTACGAGA ATCGCTGTCCTCTCCTTC | 72 |
| Bubble_2 | CTTCCTCTCCTGCGACAGACAGCTTCCATTCCTTGCC TGCTCTCTTCCCTCTC | 74 |
| Primer_224 | CGAATCGTAACCGTTCGTACGAGAATCGCT | 65 |

| | | |
|---|---|---|
| P_end_*yihO* | GCACTGGCAATTATTGCTGC | 55.5 |
| RP_Beg_y*ihO* | CCATAGTAAGCAGGCATCCC | 55.3 |
